# Feedback brings scene information to the representation of occluded image regions in area V1 of monkeys and humans

**DOI:** 10.1101/2022.11.21.517305

**Authors:** Paolo Papale, Feng Wang, A. Tyler Morgan, Xing Chen, Amparo Gilhuis, Lucy S. Petro, Lars Muckli, Pieter R. Roelfsema, Matthew W. Self

## Abstract

Neuronal activity in the primary visual cortex (V1) is driven by feedforward input from within the neurons’ receptive fields (RFs) and modulated by contextual information in regions surrounding the RF. The effect of contextual information on spiking activity occurs rapidly, and is therefore challenging to dissociate from feedforward input. To address this challenge, we recorded the spiking activity of V1 neurons in monkeys viewing either natural scenes or scenes where the information in the RF was occluded, effectively removing the feedforward input. We found that V1 neurons responded rapidly and selectively to occluded scenes. V1 responses elicited by occluded stimuli could be used to decode scene identity and could be predicted from those elicited by non-occluded images, indicating that there is overlap between visually-driven and contextual responses. We used representational similarity analysis to show that the structure of V1 representations of occluded scenes measured with electrophysiology in monkeys correlates strongly with the representations of the same scenes in humans measured with fMRI. Our results reveal that contextual influences alter the spiking activity of V1 in monkeys across large distances on a rapid time scale, carry information about scene identity and resemble those in human V1.

## Introduction

Neurons in primary visual cortex (V1) respond to specific features that are present in a small portion of visual space, known as the receptive field (RF). The feature and spatial selectivity of the RF depends on the feedforward inputs that the V1 neuron receives from the retina, via the lateral geniculate nucleus of the thalamus. The initial responses of cells in V1 are therefore largely determined by the characteristics of the stimulus within their RF. However, cells also receive numerous lateral and feedback projections that provide contextual input from the rest of the visual field^1,2^. The feedforward response is rapidly modulated by these contextual inputs, and it is therefore challenging to dissociate the influence of feedforward and contextual inputs^3^. Recent studies have used partially occluded images to reveal the influence of visual context on V1 representations in the absence of information in the RF^4–6^. If the occluder is placed over the neurons’ RF, their feedforward input is removed but the contextual inputs remain^4^.

Several neuroimaging studies have presented partially occluded images to humans, showing that V1 hemodynamic activity contains information about stimulus identity, even in the absence of feedforward information^4,7–9^. However, fMRI offers limited spatial and temporal resolution and the precise relationship between hemodynamic activity and spiking activity is not fully understood^10^. It therefore remains unknown whether the spiking activity of V1 neurons encodes contextual input in the absence of feedforward drive and how these contextual influences unfold across time. To address these questions, we presented 24 images, depicting natural (beaches, forests and mountains) and man-made scenes (buildings, highways and industry), which had been used in previous neuroimaging studies^7^, to two awake fixating macaque monkeys. The images were either partially occluded (‘occluded’ condition) or not (‘non-occluded’ condition; Figure 1A,B). We recorded multi-unit spiking activity (MUA) from a total of 175 recording sites in V1 with RFs located on the occluded region of the images and examined their responses to non-occluded and occluded images.

**Figure 1.**
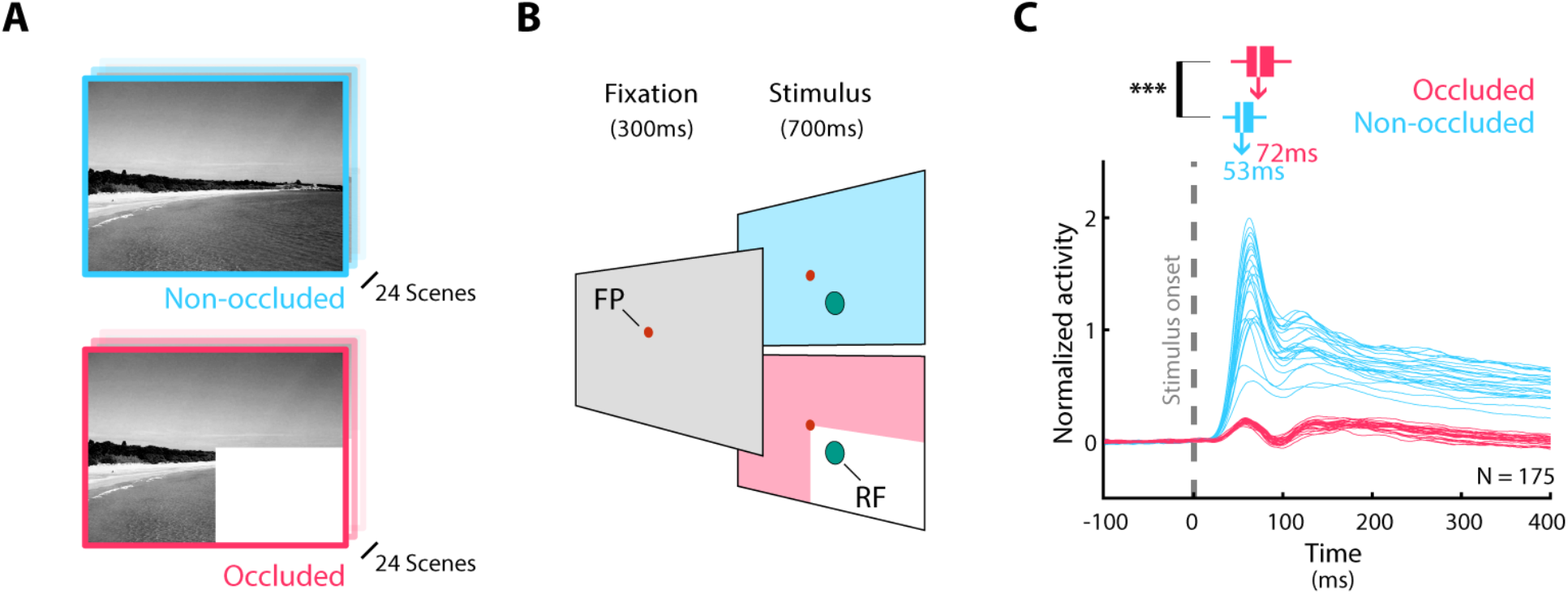
V1 response to non-occluded and partially occluded natural scenes. **a**, We presented 24 natural scenes to the monkeys; either non-occluded (top, blue) or partially occluded by a white rectangle placed in the bottom-right quadrant (bottom, pink). **b**, We recorded V1 spiking activity from two monkeys. After they maintained their gaze on a red dot (on a gray background) for 300 ms, either a non-occluded (top, blue) or partially occluded (bottom, pink) stimulus was presented for 700 ms. **c**, Mean response elicited by each of the 24 scenes (blue: non-occluded; pink: occluded condition). Traces represent the V1 population responses elicited by each scene. The latency of the response differed between the occluded and non-occluded conditions (***: p < 0.001, Wilcoxon signed rank test); bars represent the distribution of latencies of response differences between pairs of stimuli, averaged across recording sites (white line = median, box = inter-quartile range, whiskers = min/max of data).

## Results

First, we examined the mean V1 activity across recording sites elicited by each of the 24 scenes (Figure 1C). As expected, the different images elicited different activity levels in the ‘non-occluded’ condition because the information inside the neurons’ RFs varied. Neurons exhibited weaker responses in the occluded condition (p < 0.001, Wilcoxon signed rank test), and differences in activity elicited by different stimuli arose later, presumably due to extra delays associated with recurrent connections^1,2,11,12^. To evaluate the latency at which differences in response first emerged, we averaged the response to each of the 24 scenes for each recording site, before computing the 276 pairwise differences (between scene 1 and scene 2, scene 1 and scene 3, etc.). We computed the latency as the onset of their response difference (see Methods). Finally, we averaged the obtained latencies across recording sites to obtain a population estimate. For the non-occluded condition, the median latency was 53ms, which was shorter than the latency of 72ms for the occluded condition (p < 0.001, Wilcoxon signed rank test). The latency for the non-occluded stimulus was in accordance with the expected latency for the onset of V1 responses^13^, but the latency of 72ms for the contextual influences was shorter than typical latencies reported for contextual effects in e.g. figure-ground paradigms (∼100ms, e.g. ref. ^13^). We examined whether the delayed contextual modulation could be explained by a gradual propagation of activity through horizontal connections within V1, coming from nearby neurons whose RFs are not occluded^14^. However, the latency of response differences in the occluded condition did not depend on the distance between the RF and border of the occluder (Figure S1), suggesting that the contextual response arrives simultaneously across the occluded region. Such a pattern of latencies is consistent with a feedback effect from downstream visual areas, as feedback effects can span large areas of the visual scene and are therefore relatively independent of the distance from the border of the occluder^15^.

We next examined whether V1 spiking activity could be used to discriminate between the 24 scenes using a decoding approach. We trained a multi-class linear classifier to discriminate between the 24 scenes. We trained independent classifiers at each time-point (i.e. a time window of 1.3ms ±30ms because the data was sampled at 770 Hz and smoothed with a Gaussian kernel, see Methods) from 100ms before stimulus onset until 400ms thereafter. The classifier was trained and cross-validated on independent trials and we repeated the procedure both within and across the two stimulus conditions (Figure 2). As expected, training and testing on the non-occluded images yielded a high generalization accuracy (Figure 2A, blue) that was stable over time. The same analysis in the occluded condition (Figure 2A, pink) revealed that the decoding of occluded images in V1 was significant and stable over time although the average decoding accuracy was lower than for the non-occluded images (p < 0.001, Wilcoxon signed rank test)^4,7^.

**Figure 2.**
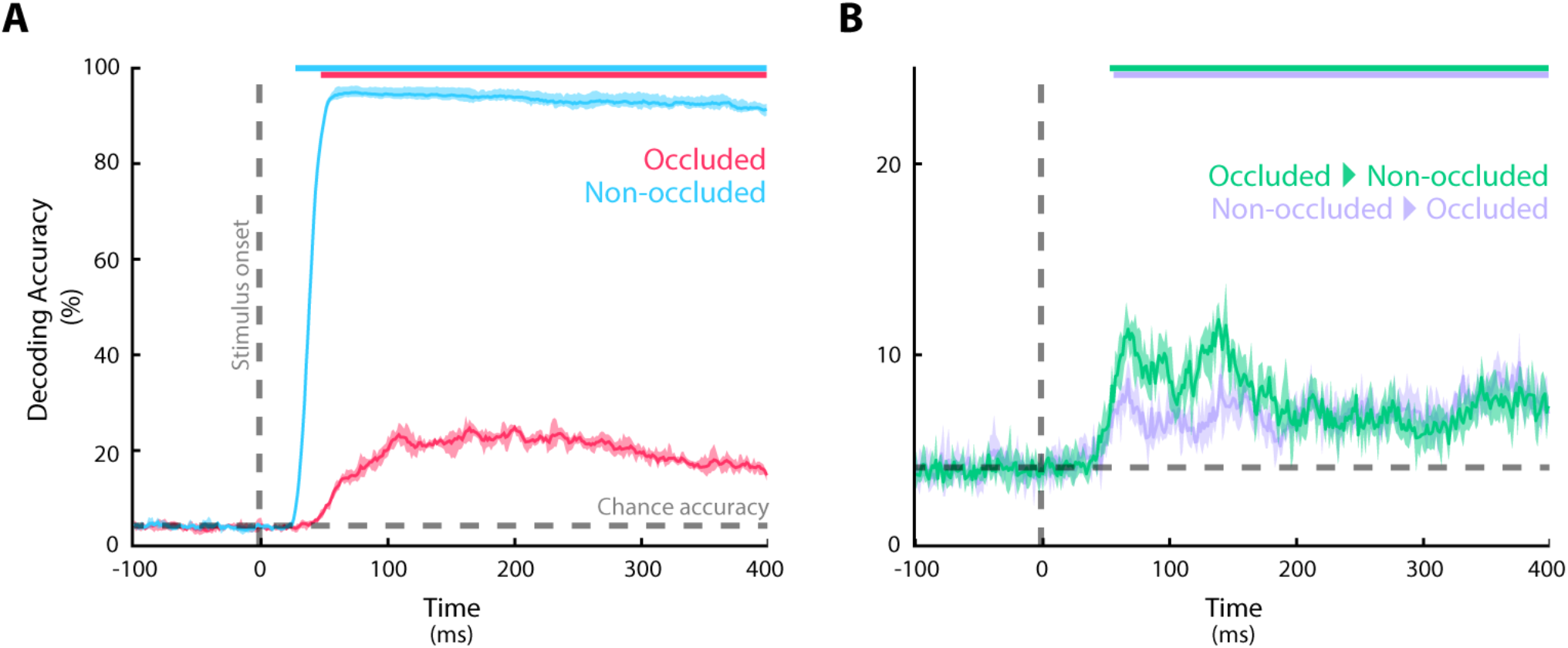
Neural decoding reveals similar contextual representations during V1 processing of non-occluded and occluded natural scenes. **a**, Results of decoding within the non-occluded (blue) and occluded (pink) conditions. The colored bars at the top depict significant time-points (p<0.05, permutation test, false discovery rate (FDR) corrected); shaded regions indicate the interval (min/max) in independent repetitions of the cross-validation procedure; the horizontal gray dashed line indicates chance level while the vertical black dashed line indicates the stimulus onset. **b**, Cross-decoding results. In purple: non-occluded_train_/occluded_test_. In green: occluded_train_/non-occluded_test_.

To examine whether the V1 representation of occluded images is similar to that of non-occluded images, we examined the cross-decoding performance. We trained a decoder on the data from the non-occluded images and tested it on data from the occluded images, and vice versa. We observed significant generalization in both directions (Figure 2B), although lower than that observed when decoding within conditions. The median accuracy was higher when training on the occluded and testing on the non-occluded condition than vice versa (p < 0.001, Wilcoxon signed rank test across time). To explore whether the code was stable or variable, we examined the temporal generality of the decoders (e.g. training at 100ms after stimulus onset and testing at 200ms, etc.) by repeating the procedure for each pair of time-points^16^. This additional analysis did reveal consistent time shifts suggesting relatively stable representations (Figure S2).

Previous studies have demonstrated that the identity of scenes can be decoded from occluded image regions in area V1 with fMRI in humans^3,4,7^. Is there a relationship between the representations of occluded image regions measured with fMRI in humans and with electrophysiology in monkeys? To investigate this question, we compared the spiking activity recorded in the two monkeys to neuroimaging data obtained from 18 human subjects who viewed the same natural scenes that were used here^7^ with a representational similarity analysis (RSA)^17,18^. For the human data, we computed dissimilarity matrices using the multi-voxel fMRI patterns in parts of human V1 that represented occluded portions of the scene (Figure 3A). Each entry in the representational dissimilarity matrix (RDM) of Figure 3A represents the similarity of the multivoxel response patterns between two pictures.

**Figure 3.**
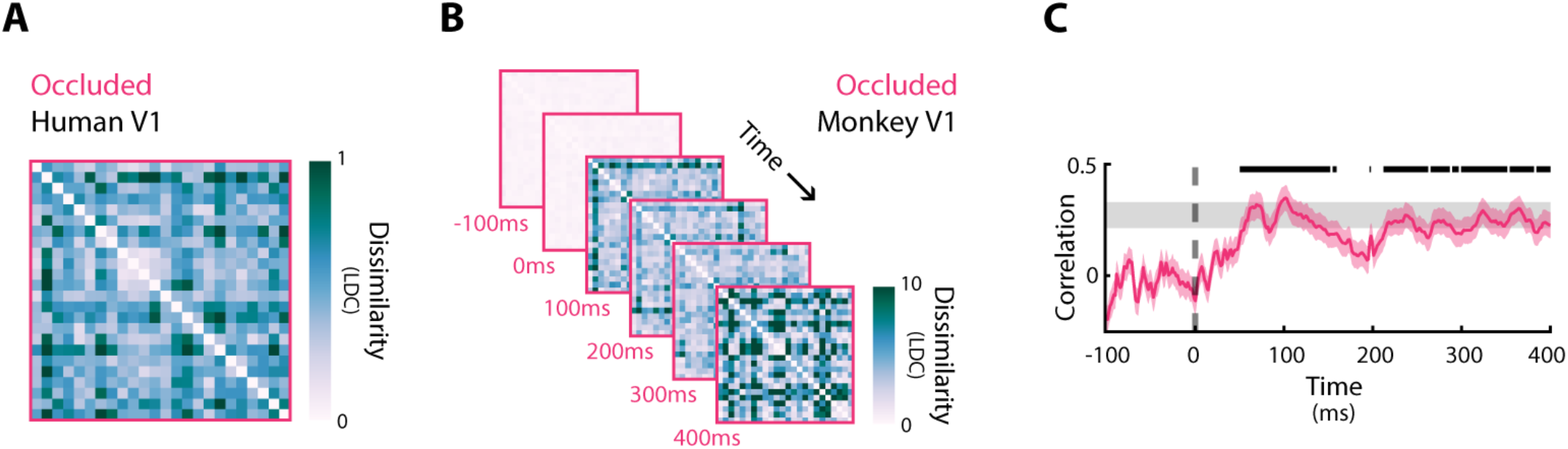
V1 representations in monkey and humans are highly correlated. **a**, Average RDM of V1 voxels across 18 subjects in the occluded region of the same images, measured with fMRI. LDC, linear discriminant contrast (see Methods). **b** RDM of monkey V1 spiking activity across time points. **c**, The Spearman’s correlation between monkey RDMs at successive time points and the human RDM. The black marker on top denotes time points with significant correlation (*p* < 0.05, permutation test, FDR corrected); the gray shaded region indicates the interval of the highest correlation that is theoretically possible given the noise in the data (i.e. noise ceiling); the shaded pink region represents bootstrapped SEM (1000 iterations).

The monkey data had a better temporal resolution than fMRI and we therefore computed monkey RDMs at successive time-points (Figure 3B)^17^. To compare the similarity between monkey and human representations of occluded image regions, we calculated the Spearman’s rank correlation between the monkey RDMs at successive time points and the human RDM (Figure 3C). We found a robust correlation that reached the noise ceiling ∼100ms after stimulus onset, meaning that the correlation between V1 representations of monkeys and the human was at least as high as that between individual human participants and their average. These results demonstrate a remarkable similarity of V1 response patterns elicited by contextual information outside the neurons’ RFs across species, using measurements from different modalities (fMRI responses in humans vs. spiking activity in monkeys).

## Discussion

Our results demonstrate strong and complex contextual influences on V1 spiking activity. We found that the activity of V1 neurons is influenced by visual scene identity, starting around 70ms after stimulus onset, even if their RFs were located on occluded regions. This latency is remarkably fast, much earlier than what has been observed in previous studies on contextual influences in V1 (e.g. refs.^13,19,20^) and is likely due to top-down rather than horizontal connections. The shorter latency of contextual influences compared to previous studies might be explained by differences between tasks: previous experiments employed a limited set of artificial geometrical stimuli, which elicit responses that differ from those elicited by natural scenes^21–23^. Previous reports also used tasks requiring spatial attention shifts^13,24^, while the current study recorded neuronal responses during passive viewing. In line with our observation, a recent study found that the saliency of contextual information modulates V1 neurons with a short latency (∼80ms) during viewing of natural images, when carefully controlling feedforward information^23^. Hence, contextual modulation of V1 neurons happens faster in natural vision than previously estimated using artificial stimuli^2,23,25^.

How is the information about scene identity that is sent back to occluded V1 representation structured? Studies in humans have suggested that this top-down signal correlates with simplified scene information such as a sketch of the visual features that might be present in the occluded region^7^. Studies in mice using occluded grating-stimuli have found evidence that feature-specific feedback signals alter spiking activity in V1^6,26^. Our cross-decoding results demonstrate overlap between the informational structure in the occluded and non-occluded signals, but with an asymmetry: decoders that were trained on visual information did not generalize as well as decoders that were trained on occluded information. This suggests that the signal that is present in occluded regions of V1 may be biased towards more abstract, high-level information rather than the exact predicted contents of the RF. We note, however, that the differences between the two decoders were relatively small and future-studies will be needed to better understand the contents of the top-down signal.

Finally, we found that V1 scene representations derived from human neuroimaging data correlate strongly with spiking activity in monkeys. Our finding is the first direct evidence of similarity in the V1 representation of the two species in the absence of feedforward information. This result represents an important step in bridging the complementary, but somewhat fragmented, lines of research in humans and non-human primates, as it suggests that what we know about the complexity of the extra-classical RF response from experiments in monkeys is also likely to generalize to humans^27^. These results lay the groundwork for future research on the exact structure of the information that is sent back to occluded regions of V1, and on its potential role in shaping the activity in V1 regions devoid of feedforward information^28,29^

## Methods

### Training of the monkeys

All procedures complied with the NIH Guide for Care and Use of Laboratory Animals and were approved by the institutional animal care and use committee of the Royal Netherlands Academy of Arts and Sciences (KNAW). Two macaque monkeys (males, 7 and 8 years old) participated in the experiments. They were socially housed in stable pairs in a specialized primate facility with natural daylight, controlled humidity and temperature. The home-cage was a large floor-to-ceiling cage which allowed natural climbing and swinging behavior. The cage had a solid floor, covered with sawdust and was enriched with toys and foraging items. Their diet consisted of monkey chow, supplemented with fresh fruit. Their access to fluid was controlled, according to a carefully designed regime for fluid uptake. During weekdays, the animals received water or diluted fruit juice in the experimental set-up upon correctly performed trials. We ensured that the animals drank sufficient fluid in the set-up and supplemented the animals with extra fluid after the recording session if they did not drink enough. During the weekend, they received a full bottle of water (700-940 ml per day) in the home cage. The animals were regularly checked by veterinary staff and animal caretakers and their weight and general appearance were recorded daily in an electronic logbook during fluid-control periods.

### Surgical details

We implanted a 3D-printed titanium head-post on the monkey’s skull under aseptic conditions and general anesthesia as reported previously^30^. The monkeys were trained to direct their gaze to a 0.2° diameter fixation dot and hold their eyes within a fixation window (1.1° diameter). They then underwent a second operation to implant arrays of micro-electrodes (Utah-probes, Blackrock Microsystems) over opercular V1 and V4 (eight 5×5 arrays in monkey B; sixteen 8×8 arrays in monkey L). For the present study we only used the recordings from the V1 arrays.

### Electrophysiology

We recorded neuronal activity from 144 recording sites in V1 in monkey B (6 arrays) and 896 in monkey L (14 arrays). Neural signals were referenced to a subdural electrode and amplified using 32 channel Tucker-Davis Technologies ZIF-clip headstage amplifiers (monkey B) or Blackrock microsystems Cereplex-M headstage amplifiers. The amplified signal was sampled at 24.4kHz using a Tucker-Davis Technologies RZ2 system (monkey B) or 30kHz using a Blackrock microsystems system (monkey L). We measured the envelope of multi-unit activity by band-pass filtering the signal offline (2nd order Butterworth filter, 500 Hz-5 KHz, filtfilt.m in MATLAB) to isolate high-frequency (spiking) activity. This signal was rectified (negative becomes positive) and low-pass filtered (corner frequency = 200 Hz) to produce the envelope of the high-frequency activity, which we refer to as MUA^31^. The MUA signal was down-sampled to 770Hz and stored for further analysis. The MUA signal reflects the population spiking of neurons within 100-150 μm of the electrode and the population responses are very similar to those obtained by pooling across single units^31,32^.

### Receptive Field Mapping

We mapped the RFs of each recording site in V1 using a drifting luminance-defined bar that moved in one of eight directions (every 45°). The response to each direction was fitted with a Gaussian function and the borders of the RFs were calculated as described previously^31^.

### Stimulus presentation

We used a dataset of 24 natural scenes and a version of the same images with the bottom-right quadrant occluded by a white rectangle (taken from ref. ^7^). Images spanned several categories: beaches, buildings, forests, highways, industry, and mountains. Stimuli were presented in two different setups. For monkey B, we used a CRT monitor with refresh rate of 75 Hz and resolution of 1024×768 pixels, viewed from a distance of 49.5 cm. The monitor provided a field-of-view of 43.5 × 32.6°. For monkey L, we used a CRT monitor with refresh rate of 85 Hz and resolution of 1024×768 pixels viewed from a distance of 50 cm. The monitor provided a field-of-view of 39.6 × 29.7°. In both setups, the eye position was recorded using a digital camera (Thomas Recording, 250-Hz framerate) and stimuli were generated in Matlab using the COGENT Graphics toolbox (developed by John Romaya at the LON at the Wellcome Department of Imaging Neuroscience) and custom control software^33^.

At the start of the trial, the monkey directed its gaze to a fixation point on a gray background (luminance 14 cd×m^−2^ for monkey B; 17.5 cd×m^−2^ for monkey L). We presented the image once the monkey had maintained fixation for 300ms, and the animal had to maintain fixation for an additional 700ms during stimulus presentation. Reward was delivered after every correct trial. Aborted trials (i.e., when the monkeys did not maintain fixation for 700ms) were repeated at the end of the recording session.

### Selection of recording sites and inclusion of data

To normalize MUA, we first subtracted the mean activity obtained during the pre-trial period, in which the animal was fixating (−200 to 0ms relative to stimulus onset) and divided activity by the maximum smoothed (25ms Gaussian kernel) peak response (0-200ms from stimulus onset)^13,23^. We only included recording sites with a sufficient signal-to-noise ratio (SNR), estimated by dividing the maximum of the initial peak response by the standard deviation of the baseline activity across trials. If the SNR of a recording site was less than 1 across recording days, we removed that recording site from the analysis. In addition, we excluded recording sites whose RF overlapped with non-occluded parts of the stimuli in the occluded condition. We only included sites whose RFs were fully inside the occluder and whose RF centers were at least 2° away from both the horizontal and vertical meridian, so that a total of 175 recording sites remained (18 in monkey B and 157 in monkey L). The mean activity, response latency and decoding accuracy were comparable across the two monkeys, thus we aggregated the data in the analyses (Figure S3). Decoding was also significant in individual monkeys for the occluded stimuli (average response after stimulus onset: both ps < 0.001, permutation test). In total, we recorded 9,600 correct trials from monkey B and 9,864 trials in monkey L.

### Analyses of latency

To compute the latency of neural responses, a function was fitted to the time-course of interest (e.g. the difference in activity elicited by different stimuli)^13,23,34^. The function was derived from the assumption that the onset of the response had a Gaussian distribution, and that a fraction of the response dissipated exponentially, yielding the following equation:

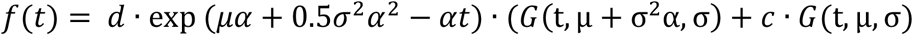

where *G*(t, μ, σ) is a cumulative Gaussian density with mean *μ*, and standard deviation *σ, α*^−1^ is the time constant of the dissipation, and *c* and *d* represent the contribution of the non-dissipating and dissipating components. The function was fit using non-linear least squares (fit.m in MATLAB). The latency was defined as the point at which the fitted function reached 33% of its maximum.

### Neural decoding

All trials were used for decoding, after Gaussian smoothing (*σ* = 25 *ms*). Training and generalization were performed using data from a single time point with a multiclass (24 classes) linear discriminant analysis classifier, implemented in Matlab (the *fitcdiscr* function), according to the following equation:

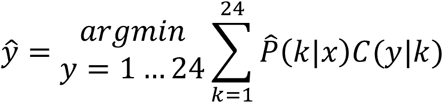

where *ŷ* is the predicted classification, 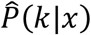 is the posterior probability of class *k* for observation *x* and *C*(*y*|*k*) is the cost of classifying an observation as *y* when the true class is *k*. In addition, we also repeated the procedure for every possible pair of time points between training and generalization - a procedure known as temporal generalization^16^ (Figure S2).

V1 responses elicited by non-occluded stimuli had a higher SNR than those elicited by the occluded condition (measured as the ratio between the mean and standard deviation of the response). It has been shown that training on a dataset with higher SNR leads to poor generalization when testing on a dataset with lower SNR^35^. In order to address this issue, when generalizing from the non-occluded to the occluded condition, we conducted extra trial averaging (10 random trials, constant across cross-validation folds) in the test-set of occluded data to match the SNR of V1 responses elicited by the non-occluded stimuli (SNR after stimulus onset was 3.1 for the non-occluded condition, 1.8 for the occluded condition and 3.2 for the matched occluded condition).

To avoid overfitting and to obtain a robust estimate of the generalization performance, we repeated a cross-validation procedure five times (i.e. we trained the decoders on a randomly selected 80% of the trials and tested them on the held-back 20% of trials and repeated this procedure 5 times) and averaged across the accuracies. We estimated the statistical significance of decoding with a permutation test. First, we built a permuted null-distribution by repeating the same procedure 100 times and shuffling the labels of the stimuli (keeping the time points for training and generalization the same). Second, we selected the upper tail (highest 30%) of the null distribution, and we fitted a Pareto function to the tail to estimate extreme values in full^36^. Third, we identified the percentile corresponding to each accuracy value in the null-distribution. Finally, where appropriate, we corrected for multiple comparisons using the false discovery rate (FDR) correction^37^.

### Correlation analysis

We used RSA to gauge the similarity between monkey V1 spiking activity and human V1 functional MRI responses, using data from 18 human subjects (see ref.^7^ for details on MRI acquisition and preprocessing). For both monkey and human V1, we computed a RDM based on pairwise distances, computed as linear discriminant contrast (LDC, i.e. the cross-validated Mahalanobis distance between either voxel activations (for humans) or recording sites (for monkeys) activations to different stimuli)^38^, and we used the same cross-validation scheme discussed above. For human V1, only voxels representing the occluded image region were included, as reported in ref. ^7^. To calculate the similarity between each time point in the data of monkey V1 and the average dissimilarity matrix across human subjects, we used Spearman’s rank correlation. To compute the statistical significance of each correlation value, we employed a permutation test: for each time point in the monkey V1 data, we built a null-distribution repeating the same procedure while randomly shuffling the order of elements in the monkey V1 dissimilarity matrix a thousand times. Then, we computed the p-value of each real correlation value by identifying the percentile corresponding to its value in the null-distribution^39^. Finally, we corrected for multiple comparisons (across time-points) using an FDR correction^37,39^. In addition, we computed the interval of the highest correlation that was theoretically possible, given the noise in the data (i.e., noise ceiling), using the procedure described in ref.^40^.

## Acknowledgments

We thank Kor Brandsma, Anneke Ditewig and Lex Beekman for biotechnical support, and Rick Schuurman for assistance during surgeries. We thank Blackrock Microsystems for technical assistance and collaboration. Funding: The work was supported by NWO (STW-Perspectief grant P15-42 “NESTOR” and Crossover grant 17619 “INTENSE”), a European Union FP7 ERC grant (339490 “Cortic_al_gorithms”), the European Union Horizon 2020 Framework Program under specific grant agreement 785907 and 945539 “Human Brain Project” SGA2 and SGA3, and the Biotechnology and Biological Sciences Research Council (BBSRC) ‘Layer-specific cortical feedback’.

## Author contributions

P.P., M.W.S and P.R.R conceived of the study. F.W., X.C., M.W.S., and A.G. performed the experiments. P.P. analyzed data with contributions from F.W., A.T.M. and M.W.S. P.R.R. and L.M. (with L.S.P.) obtained funding. A.T.M., L.S.P. and L.M. provided data for the study. M.W.S and P.R.R oversaw all aspects of the project. P.P., P.R.R. and M.W.S wrote the paper with contributions from all authors.

