## Supplementary figures for "Feedback brings scene information to the representation of occluded image regions in area V1 of monkeys and humans"

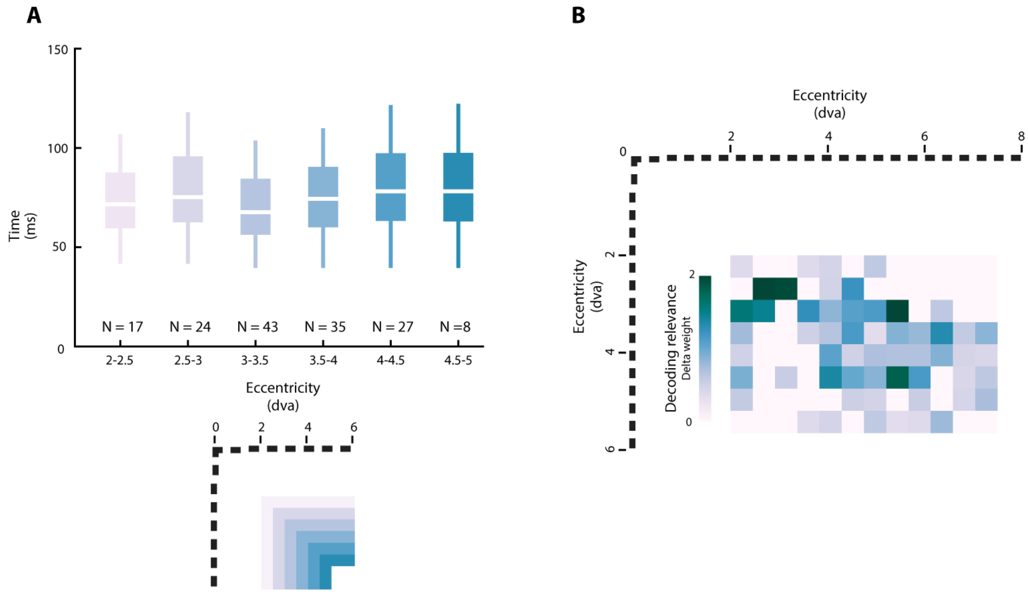


**Figure S1. Absence of an influence of the distance of RFs to the image borders on stimulus discrimination latency and decoding accuracy.**

**A.** The latency of the differences between the responses obtained for pairs of stimuli in the occluded condition did not depend on the distance between the RFs and the non-occluded portion of the image ($p > 0.05$, Kruskal-Wallis test). Top: binned recording sites (0.5 degrees of visual angle [dva] width) of monkey L according to the distance between their RF and the borders of the non-occluded image region (bottom). **B.** Decoding accuracy did not depend on the distance between the RFs and the non-occluded image region (x-axis: horizontal meridian; y-axis: vertical meridian). Recording sites were binned into an 8x12 grid on the basis of their RF position. Colors represent the average influence of recording sites on decoding within each bin (0.5 x 0.5 dva). Influence was estimated as the weight of recording sites in the decoder.

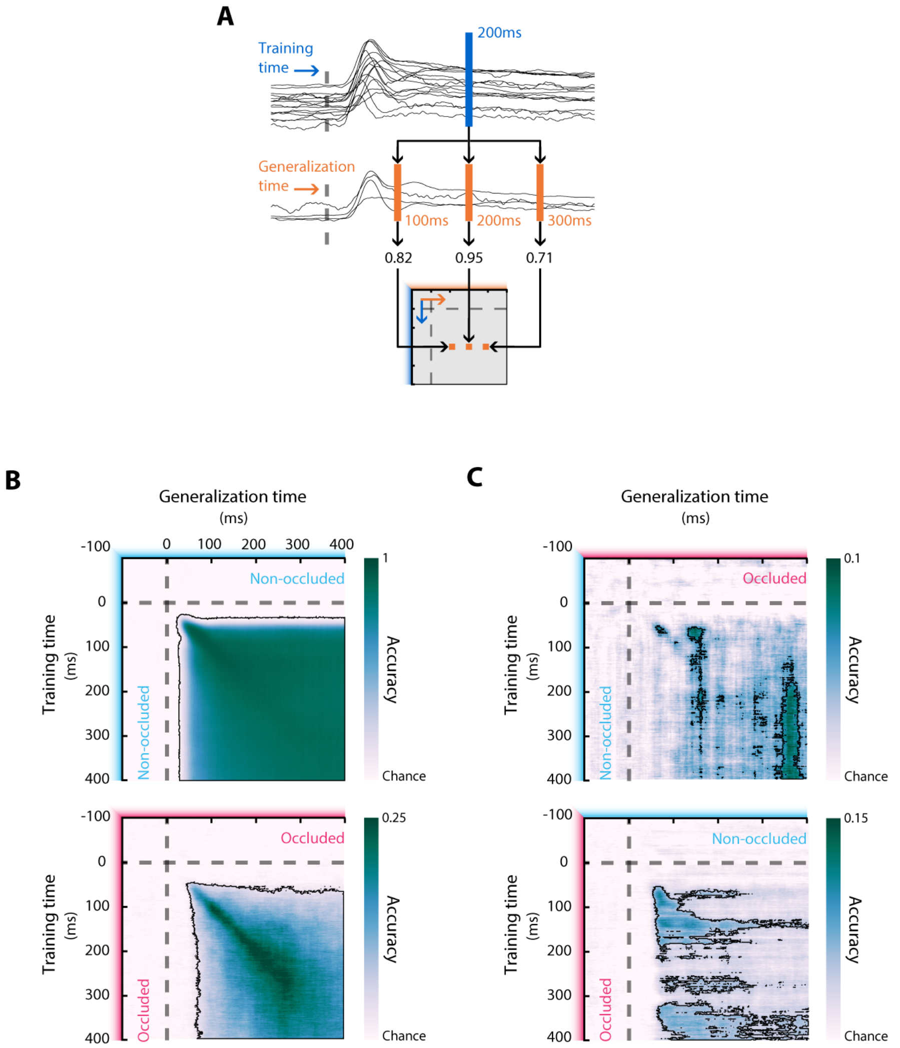


**Figure S2. Time-generalized decoding.**

**A.** Similarly to the decoding analysis of the main results section, we trained a linear classifier and assessed how decoding generalized to different time-points, from 100ms before stimulus onset until 400ms thereafter. We constructed generalization matrices within (**B**) and across (**C**) occluded and non-occluded conditions.

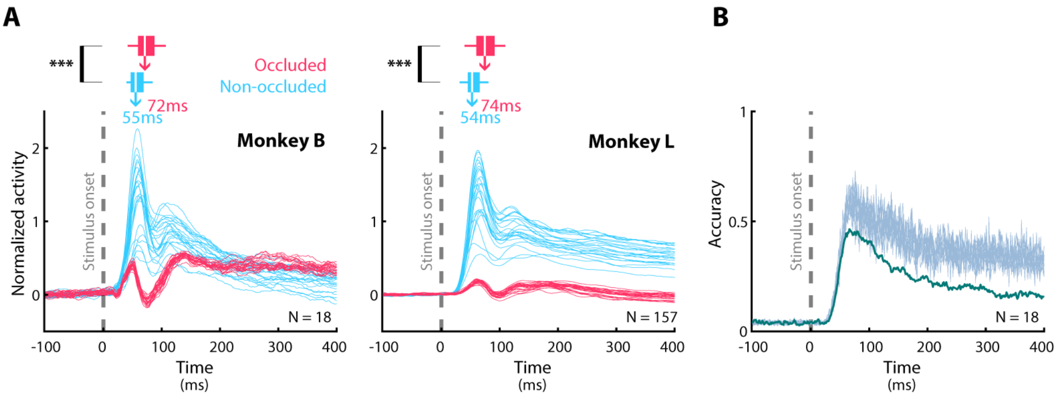


**Figure S3. Mean responses, latencies and decoding accuracy are comparable between monkeys.**

**A.** Mean responses and discrimination latencies for monkey B (left) and monkey L (right). The responses were similar and the difference in the delay in the discrimination latency between occluded vs. non-occluded stimuli was similar in the two animals (***: $p < 0.001$, Wilcoxon signed rank test). **B.** When we equated the number of recording sites in the two monkeys (sampling 10 different random combinations of 18 channels in monkey L), decoding accuracy of non-occluded stimuli in monkey B (green) was comparable to that in monkey L (gray).
